# Elevated sleep need in a stress-resilient *Drosophila* species

**DOI:** 10.1101/2023.05.27.542279

**Authors:** Jessica Yano, Ceazar Nave, Katherine Larratt, Phia Honey, Cassandra Jingco, Makayla Roberts, Damion Trotter, Xin He, Gazmend Elezi, Julian P. Whitelegge, Sara Wasserman, Jeffrey M. Donlea

## Abstract

Sleep is broadly conserved across the animal kingdom, but can vary widely between species. It is currently unclear which types of selective pressures and sleep regulatory mechanisms influence differences in sleep between species. The fruit fly *Drosophila melanogaster* has become a successful model system for examining sleep regulation and function, but little is known about the sleep patterns and need for sleep in many related fly species. Here, we find that *Drosophila mojavensis*, a fly species that has adapted to extreme desert environments, exhibits strong increases in sleep compared to *D. melanogaster.* Long-sleeping *D. mojavensis* show intact sleep homeostasis, indicating that these flies carry an elevated need for sleep. In addition, *D. mojavensis* exhibit altered abundance or distribution of several sleep/wake related neuromodulators and neuropeptides that are consistent with their reduced locomotor activity, and increased sleep. Finally, we find that in a nutrient-deprived environment, the sleep responses of individual *D. mojavensis* are correlated with their survival time. Our results demonstrate that *D. mojavensis* is a novel model for studying organisms with high sleep need, and for exploring sleep strategies that provide resilience in extreme environments.

## Results and Discussion

Sleep is widely conserved across the animal kingdom, but the amount of time that individuals spend asleep varies widely among species. While previous studies and meta-analyses have examined physical and life history traits that correlate with interspecies variations in sleep ^1–7^, the feasibility of systematic comparisons of sleep across related vertebrate species is limited. In contrast, the *Drosophila* genus provides a diverse range of species, including the genetic model species *D. melanogaster,* many of which can be cultured and behaviorally monitored in standard laboratory conditions^8^. To begin sampling the sleep strategies of *Drosophila* species, we compared sleep in *D. melanogaster* and in the cactophilic species *D. mojavensis*. We show that *D. mojavensis* exhibits increased sleep time across the day and night compared to *D. melanogaster*, and that desert-adapted *D. mojavensis* flies respond to sleep loss with a homeostatic increase in sleep drive. We observe several changes in sleep- or wake-related neuromodulator distribution: long-sleeping *D. mojavensis* flies exhibit high levels of serotonin, decreased abundance of wake-promoting octopamine, and reduced numbers of cells expressing the circadian output peptide Pigment Dispersing Factor (PDF). Finally, we examine contributions of elevated sleep to stress resilience in *D. mojavensis* by measuring starvation and dehydration responses. Long-sleeping *D. mojavensis* flies exhibit extended survival during food or food and water deprivation compared to *D. melanogaster*, and individual sleep time of *D. mojavensis* correlates positively with survival time while flies are starved and dehydrated. These results indicate that *D. mojavensis* exhibits an increased need for sleep relative to *D. melanogaster*, and that adaptations in sleep may contribute to increased stress resilience in desert-adapted flies.

*Drosophila melanogaster* has become a popular genetic model system to study sleep and circadian rhythms^9–11^. While focus on this model species permits the rapid development and proliferation of genetic tools and mechanistic frameworks, few studies have examined sleep in related species that are adapted to thrive in a variety of environmental conditions. Increased sleep is a behavioral adaptation that is hypothesized to support resistance to nutrient scarcity^12^, and artificial selection for starvation resistance in *D. melanogaster* can result in increased sleep time^13^. To test whether similar changes in sleep strategies might correlate with interspecific changes in stress resistance, we compared sleep and starvation/dehydration responses in *D. melanogaster* and the cactophilic species *D. mojavensis* (See **Fig. 1a** for phylogenetic tree, based on ^14^)*. D. mojavensis* are found in desert regions of Mexico and the southwestern USA and includes four geographically segregated subspecies: *D. moj. mojavensis*, *D. moj. baja*, *D. moj. sonorensis*, and *D. moj. wrigleyi* from the Mojave Desert, Baja California, Sonoran Desert, and Santa Catalina Island, respectively ^15–17^. We measured sleep in all four *D. mojavensis* subspecies and in two wild-type stocks of *D. melanogaster* (*Cs*^18^ and *Pcf*^19^) using multibeam *Drosophila* activity monitors. Each *D. mojavensis* subspecies exhibits significantly elevated sleep throughout the day and night compared to *D. melanogaster* **(Figs. 1b-c)**. To test whether hypersomnolence in *D. mojavensis* can be attributed to an elevated pressure to maintain sleep and/or to an increased drive to initiate sleep episodes, we quantified the likelihood that a sleeping fly would awaken (P(wake); **Fig. 1d**) or that a waking fly would fall asleep (P(doze); **Fig. 1e**)^20^. Each of the four *D. mojavensis* subspecies exhibits reduced P(wake) and elevated P(doze) compared to *D. melanogaster*, consistent with both strengthened sleep maintenance and an elevated pressure to fall asleep. Along with increased sleep time, *D. mojavensis* also exhibits reduced waking locomotor activity **(Fig. 1f)**, consistent with previous reports^21^. To test for variations in sleep across days, we measured locomotion in *D. moj. mojavensis* flies across a 7-day period and found that daily sleep varies between individuals, but remains stable over time for single flies **(Fig. 1g-h)**.

**Figure 1.**
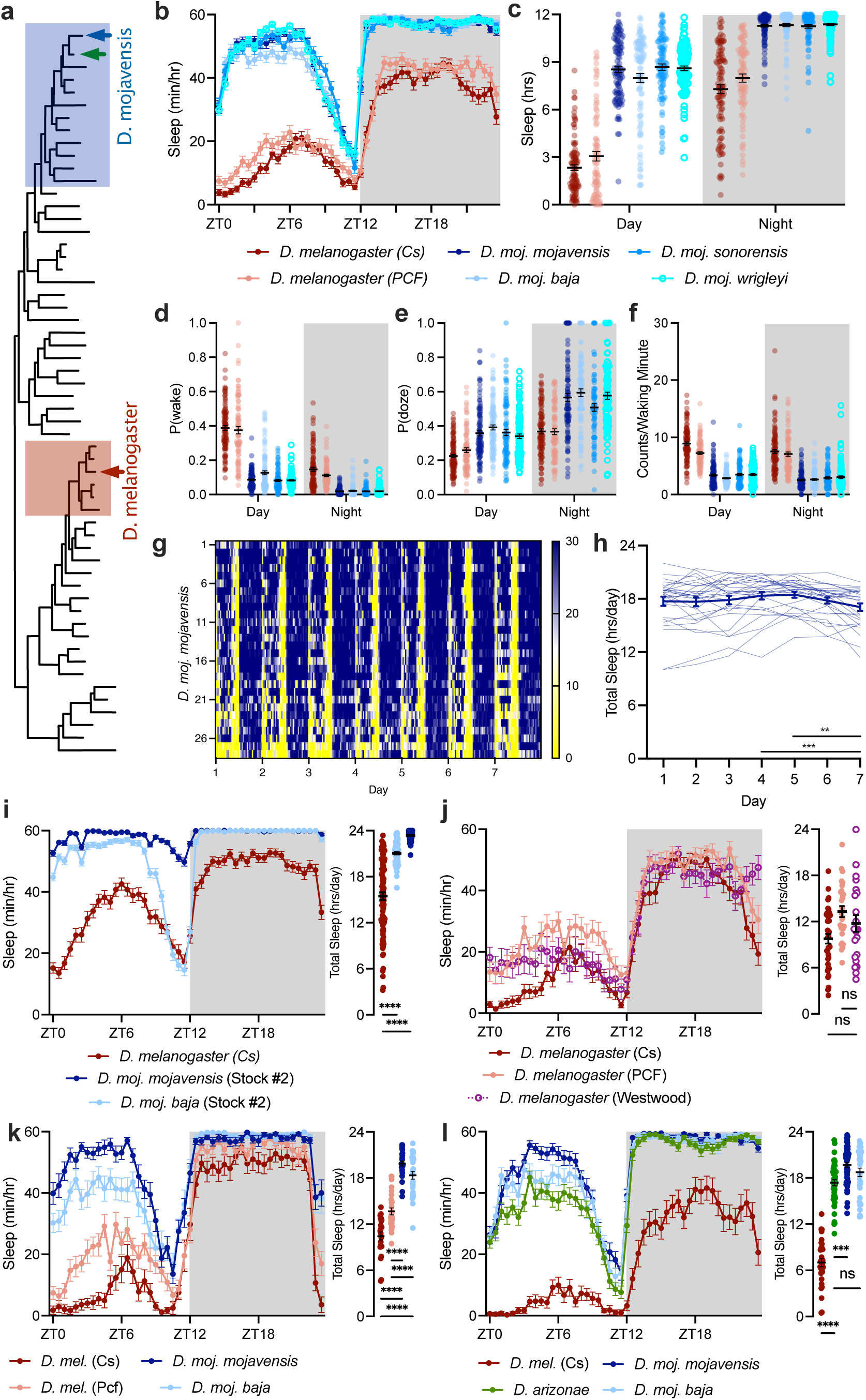
Elevated sleep time in desert-dwelling *Drosophila mojavensis*. **(a)** Phylogenetic tree of *Drosophila* species, *D. melanogaster* (red) and *D. mojavensis* (blue) are marked with arrows. Based on ^14^. Shading represents Melanogaster (red) and Repleta (blue) species groups that include *D. melanogaster* and *D. mojavensis,* respectively. **(b)** 24h sleep timecourse for wild-type *D. melanogaster (Cs,* dark red; *Pcf*, light red) and four subspecies of *D. mojavensis* (blue). Two-way repeated measures ANOVA finds a significant genotype-by-time interaction (F_(235,27213)_=16.99, p<0.0001). **(c)** Day and night sleep totals for *D. melanogaster* (*Cs*, dark red*; Pcf,* light red) and *D. mojavensis* (blues). Two-way repeated measures ANOVA finds a significant genotype-by-time interaction (F_(5,577)_=24.981, p<0.0001). **(d-e)** P(wake) **(c)** and P(doze) **(d)** during the day and night for *D. melanogaster* (reds) and *D. mojavensis* (blues) stocks. Two-way repeated measures ANOVA detects a significant genotype-by-time interaction for P(wake) (F_(5,579)_=75.43, p<0.0001) and for P(doze) (F_(5,553)_=5.628, p<0.0001). **(f)** Waking activity (position movements/waking minute) is decreased in *D. mojavensis* subspecies (blues) relative to *D. melanogaster* (reds). Two-way repeated measures ANOVA finds a significant main effect of genotype (F_(5,576)_=139.4, p<0.0001). For panels **(a-f)**, n= 101 *Cs,* 82 *Pcf*, 100 *D. moj. moj.*, 100 *D. moj. baja*, 93 *D. moj. sonorensis*, 106 *D. moj. wrigleyi*. **(g-h)** Sleep timecourse heatmap **(g)** and daily sleep totals **(h)** for *D. moj. moj.* female flies across a 7-day experiment (n=28 flies). **(i)** Sleep timecourse (left) and total daily sleep (right) for *Canton-S* (red) and additional stocks of *D. moj. moj.* (dark blue) and *D. moj. baja* (light blue). ANOVAs find a significant genotype-by-time interaction in sleep timecourse (F_(94,13207)_=40.94, p<0.0001) and main effect of genotype for total daily sleep (F_(2, 281)_=178.9, p<0.0001, n= 96 *Canton-*S, 96 *D. moj. moj.,* and 92 *D. moj. baja*). **(j)** Sleep timecourse (left) and total daily sleep (right) for *Canton-S* (dark red), *Pcf* (light red), and flies descended from *D. melanogaster* caught in Westwood, Los Angeles (open purple circles). ANOVAs detect a significant genotype-by-time interaction in sleep timecourse (F_(94,3995)_=2.385, p<0.0001) and main effect of genotype for total daily sleep (F_(2,85)_=5.793, p=0.0044, n= 38 *Canton-S,* 28 *Pcf*, and 22 wild-caught flies). **(k)** Sleep timecourse (left) and total daily sleep (right) for *D. melanogaster* stocks (red) and two *D. mojavensis* subspecies (blue) from flies reared on Banana-Opuntia media. ANOVAs detect significant genotype-by-time interaction for the sleep timecourse (F_(141,4841)_=8.838, p<0.0001) and a significant effect of genotype for total daily sleep (F_(3, 103)_=91.08, p<0.0001, n= 24 *Canton-S*, 27 *Pcf*, 28 *D. moj. moj.,* and 28 *D. moj. baja*). **(l)** Sleep timecourse (left) and total sleep (right) for *D. melanogaster* (red), *D. mojavensis* (blues), and *D. arizonae* (green). ANOVA tests find significant genotype-by-time interaction for sleep timecourse (F_(141,7614)_=6.534, p<0.0001) and a significant effect of genotype for total sleep (F_(3,162)_=148.3, p<0.0001, n= 31 *Canton-S*, 53 *D. moj. moj*, 35 *D. moj. baja*, and 47 *D. arizonae*).

Because the *D. mojavensis* stocks that we describe above were derived from wild populations more recently than either of our wild-type *D. melanogaster* stocks, we next tested whether fly lines isolated from the wild might sleep more than those reared in lab conditions for longer periods of time. First, we found that independent stocks of *D. moj. mojavensis* and *D. moj. baja* also showed a strong increase in sleep time compared to wild-type *D. melanogaster* **(Fig. 1i)**. Next, we examined sleep in a *D. melanogaster* stock that originated from flies collected in the Westwood area of Los Angeles in 2020. *D. melanogaster* that descended from flies caught in Westwood, Los Angeles, CA showed comparable sleep amounts to *Cs* and *Pcf* laboratory strains **(Fig. 1j)**. To test the impact of diet on sleep in *D. mojavensis*, we collected freshly eclosed flies and reared them on media that included extract of opuntia cactus, a natural host for desert-adapted *D. mojavensis.* Sleep in *D. mojavensis* remained elevated relative to *D. melanogaster* when both species were fed a banana-cactus diet **(Fig. 1k)**. In addition to *D. mojavensis*, several other closely related fly species, including *D. arizonae,* also localize to deserts of the Southwest USA and Mexico and feed on cactus hosts^22^ (Denoted in **Fig. 1a** by green arrow). As shown in **Fig. 1l**, *D. arizonae* show comparable amounts of sleep as *D. mojavensis*, suggesting that elevated sleep is not exclusive to *D. mojavensis* and could be conserved across related fly species in the Repleta species group that localize to desert regions^23^.

Elevated sleep in a desert-adapted species could indicate at least two possibilities: first, that sleep provides a period of adaptive inactivity during which animals can store metabolic resources and avoid predation^5^, or alternatively, that this species has adapted an elevated need for basic functions that are fulfilled by sleep. To test whether desert-adapted *D. mojavensis* maintain an elevated sleep need, we tested whether they respond to mechanical sleep deprivation with a homeostatic rebound to recover lost sleep. Vortex stimuli delivered for 3s each minute were sufficient to strongly suppress sleep in *D. moj. mojavensis* **(Fig. 2a)** and in *D. moj. baja* **(Fig. 2b)**. Following sleep deprivation, both *D. mojavensis* subspecies showed a recovery period of significantly increased sleep compared to baseline and regained approximately 20-40% of their lost sleep after 24 hours **(Fig. 2c)**. In the 24h following deprivation, P(wake) is decreased during daytime sleep on the first recovery day after deprivation, an indication of increased sleep depth **(Extended Data Fig. 1a-b)**. Additionally, there was no decrease in locomotor activity per time awake **(Extended Data Figure 1c-d)**, indicating that waking locomotor activity was unimpaired by mechanical sleep deprivation. Following the first 24h of recovery, *D. moj. mojavensis* flies reduced their sleep nearly to baseline levels on the second recovery day **(Extended Data Fig. 1e-f)**. After finding that *D. mojavensis* exhibited a rebound in sleep time after overnight sleep loss, we next probed arousability to test for additional markers of increased sleep depth in recently deprived *D. moj. mojavensis.* Flies were either left undisturbed, sleep-deprived for 12h overnight (SD), or sleep-deprived and permitted 24h of recovery (SD+24h) before they were exposed hourly to 60s pulses of blue light. Light pulses were less likely to awaken sleep-deprived flies than rested controls; arousability returned to control levels in SD+24h flies **(Fig. 2d)**. After each light pulse, *D. moj. mojavensis* flies in the SD group had a reduced latency to fall back asleep compared to both the control and SD+24h groups **(Fig. 2e)**. These results indicate that long-sleeping *D. mojavensis* responds to mechanical sleep loss with homeostatic increases both in sleep time and intensity, consistent with the hypothesis that *D. mojavensis* have adapted an increased need for sleep.

**Figure 2.**
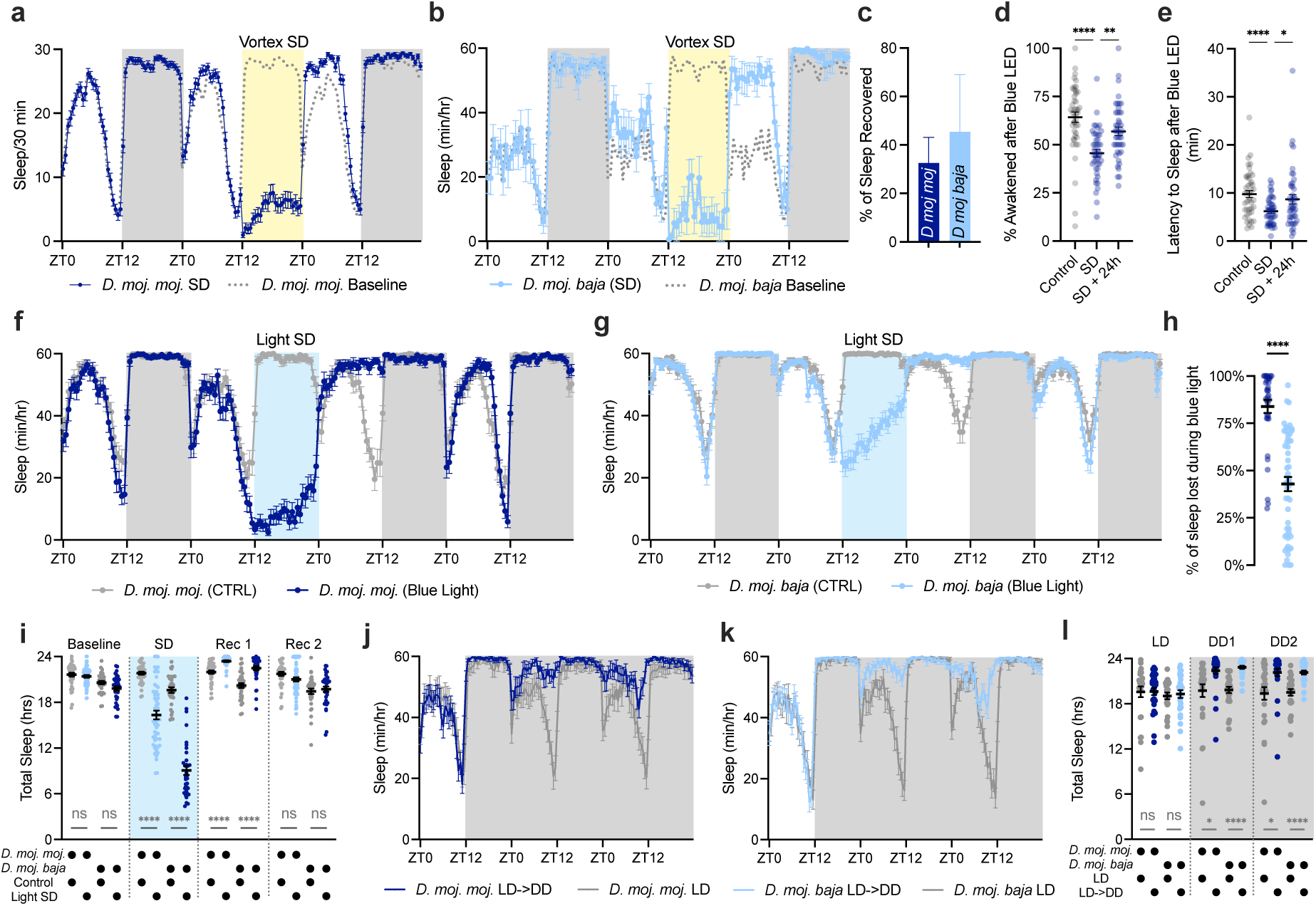
Homeostatic regulation of sleep and arousability in *Drosophila mojavensis*. **(a-b)** Sleep timecourse of *D. moj. moj.* **(a)** and *D. moj. baja* **(b)** across baseline, overnight mechanical sleep deprivation, and recovery days. Yellow shading indicates time of sleep deprivation. Dotted gray lines show baseline sleep patterns replotted on deprivation and recovery days for visual comparison (n= 77 flies in **a**, 38 in **b**). **(c)** Percentage of sleep recovered within 24h of recovery from mechanical sleep deprivation. *D. moj. moj.* shown in dark blue, *D. moj. baja* in light blue. (n= 77 for *D. moj. moj.* and 38 for *D. moj. baja* and flies/group). **(d)** Portion of sleeping *D. moj. mojavensis* flies awakened by 60s pulses of blue light. Individual data points represent group mean response rate from individual hourly light exposure trials. One-way repeated measures ANOVA finds a significant effect of condition (F_(1.874, 80.56)_=15.41, p<0.0001, n=44 trials/group). **(e)** Mean sleep latency of *D. moj. mojavensis* flies after hourly 60s pulses of blue light is reduced after mechanical sleep deprivation. Individual data points represent group mean sleep latency after individual hourly light exposure trials. One-way repeated measures ANOVA finds a significant effect of condition (F_(1.730,74.40)_=7.342, p=0.002, n=44 trials/group). **(f-g)** Sleep timecourse of *D. moj. mojavensis* **(f)** and *D. moj. baja* females **(g)** during baseline, overnight blue light exposure, and two recovery days. Blue shading shows the time of overnight light stimulation. Gray traces represent undisturbed controls and blues depict sleep for flies exposed to blue light from ZT12-24 on day 2. Two-way repeated measures ANOVAs find significant time-by-condition interactions for **(f)** (F_(191,12606)_=33.98, p<0.0001, n=33 control, 35 Light SD) and for **(g)** (F_(191,16999)_=15.98, p<0.0001, n=38 control, 53 Light SD). **(h)** Percentage of sleep lost during 12h of overnight blue light exposure in *D. moj. moj.* (dark blue) and *D. moj. baja* (light blue). Unpaired T-test t=7.593, df=86, p<0.0001, n=35 *D. moj. moj.*, 53 *D. moj. baja*. **(i)** Daily sleep totals for groups shown in **(f)** and **(g)**. Two-way repeated measures ANOVA finds a group-by-day interaction (F_(9,465)_=97.60, p<0.0001, n=33 Control *D. moj. moj.*, 35 Light SD *D. moj. moj.*, 38 Control *D. moj. baja*, 53 Light SD *D. moj. baja*). **(j-k)** Sleep timecourses for *D. moj. moj.* **(j)** and *D. moj. baja* **(k)** during one day of 12h:12h light-dark followed by two days in constant darkness. Gray traces show controls that remain on 12h:12h LD schedule, groups transferred to darkness depicted in blues. Two-way ANOVAs find significant group-by-time interactions for **(j)** (F_(143, 7436)_=6.694, p<0.0001, n=26 LD, 28 LD->DD flies/group) and **(k)** (F_(143, 7293)_=10.40, p<0.0001, n= 25 LD, 28 LD->DD flies/group). **(l)** Total daily sleep for groups shown in **(j)** and **(k)**. Two-way repeated measures ANOVA finds a significant group-by-day interaction (F_(6, 206)_=12.02, p<0.0001, n= 26 *D. moj. moj.* LD, 28 *D. moj. moj.* LD->DD, 25 LD *D. moj. baja*, 28 LD->DD *D. moj. baja*).

To further probe responses of *D. mojavensis* to acute sleep loss, we also exposed *D. moj. mojavensis* and *D. moj. baja* flies to arousing blue light for 12h overnight (ZT12-0). Overnight blue light disrupted sleep in both desert subspecies and was followed by prolonged rebound during the first recovery day **(Fig. 2f-i)**. During light stimulation, *D. moj. mojavensis* lost 83.90±3.50% (mean ± SEM, n=35) of their sleep while *D. moj. baja* reduced their sleep by 42.89±3.74% (mean ± SEM, n=53) **(Fig. 2h)**. Given that overnight light exposure significantly disrupted sleep, we next tested whether acute visual input bidirectionally influences sleep by housing *D. mojavensis* in two days of constant darkness. Both *D. moj. mojavensis* **(Fig. 2j)** and *D. moj. baja* **(Fig. 2k)** significantly increased their sleep when transferred to constant darkness after entrainment in a 12h:12h light-dark schedule. We found that in the absence of day-night light signals, the immediate increase in subjective daytime sleep persists across at least two days **(Fig. 2l)**. Previous observations of *D. melanogaster* have found either reduced or unchanged sleep when flies were housed in constant darkness ^24–27^, indicating that light-dependent modulation of sleep could be a target of evolutionary adaptation.

Research over the past 20 years identified several neuromodulators and neuropeptides that influence sleep/wake regulation in *D. melanogaster*^27–32^, but interspecies variation of these signals across fly species is not well-studied. In particular, we hypothesized that hypersomnia in *D. mojavensis* may be correlated with an upregulation of sleep-promoting signals and a decrease in arousal pathways. To identify relevant neuromodulators, we conducted liquid chromatography-mass spectrometry (LC-MS) assays of fly heads from both *D. melanogaster* and *D. mojavensis*. We found that long-sleeping *D. mojavensis* flies from all four subspecies contain a significant increase in serotonin (5-HT) and decrease of octopamine (OA) **(Fig. 3a-b)**. No uniform change in dopamine (DA) or histamine (HA) was measured between species **(Fig. 3c-d)**. 5-HT signaling promotes sleep in *D. melanogaster* ^29, 33–35^ and in vertebrates ^36–38^, while OA, a paralog of norepinephrine ^39^, drives arousal ^30, 40^. To examine whether the distribution of other wake-promoting signals might differ between these two fly species, we performed immunostaining for the arousing circadian output peptide Pigment Dispersing Factor (PDF)^41^. While *D. melanogaster* brains contain eight PDF-positive neurons in each hemisphere, four s-LNvs and four l-LNvs **(Fig. 3e)**, careful analysis reveals inconsistent PDF-expression patterns between *D. melanogaster* and *D. mojavensis* **(Fig. 3f-g)**. Specifically, *D. mojavensis* retained three to four PDF-positive l-LNvs, but showed no s-LNv cell bodies or dorsal protocerebrum projections that were labelled with anti-PDF **(Fig. 3f-g, i-j)**. A loss of PDF-immunostaining in s-LNvs has also been reported in other *Drosophila* species, indicating that selective pressures may drive reconfiguration of clock circuits as species adapt to different environments^42–45^. Together, these results indicate that elevated sleep of desert-adapted *D. mojavensis* correlates with both an increase in sleep-promoting 5-HT and reductions of arousing OA and PDF. To functionally test the role of interspecies variation in neuromodulators, we microinjected *D. moj. baja* females with 18.4 nL of either 20mM OA or vehicle control. During the first 24h after OA injections, we found that *D. moj. baja* females showed reduced sleep **(Fig. 3k-l)** and increased locomotor activity (Fig. 3m) compared to vehicle-treated siblings.

**Figure 3.**
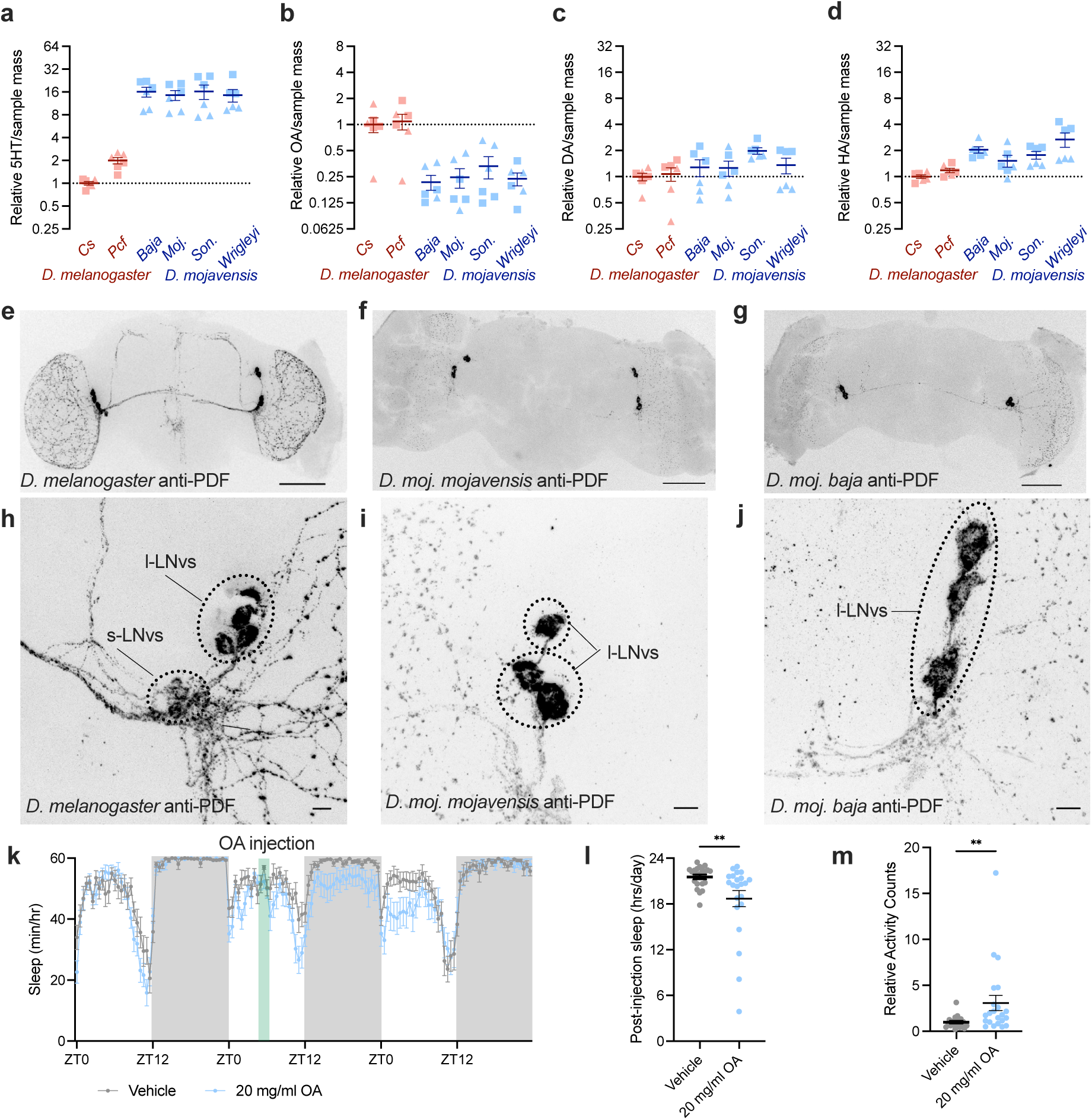
Interspecies variation of sleep- and wake-regulatory modulators between *D. melanogaster* and *D. mojavensis*. **(a-d)** Relative LC-MS/MS quantification of 5-HT **(a)**, octopamine **(b)**, dopamine **(c)**, and histamine **(d)** in heads of *D. melanogaster* wild-type stocks (reds) and *D. mojavensis* subspecies (blues). Data represent two independent experiments, each with three biological replicates per group (n=∼100 heads/biological replicate; squares represent data from Expt #1, triangles are from Expt #2). One-way ANOVAs find a significant effect of genotype for 5-HT (F_(5,30)_=10.26, p<0.0001), octopamine (F_(5,30)_=9.488, p<0.0001), and histamine (F_(5,30)_=5.950, p=0.0006), but no significant effect of genotype for dopamine (F_(5,30)_=2.465, p=0.055). **(e-g)** Immunostaining for PDF in wild-type Cs *D. melanogaster* **(e)**, *D. moj. mojavensis* **(f)**, and *D. moj. baja* **(g)**. Scale bars = 100μm. **(h-j)** PDF immunostaining of lateral-ventral neurons at 63x magnification in *D. melanogaster* **(h)**, D. moj. mojavensis **(i)**, and D. moj baja **(j)**. Dotted lines indicate the immuno-detected LNv subtypes (large or small LNv). Scale bars = 10μm. **(k)** Sleep time course for *D. moj. baja* flies that were microinjected with 18.4 nL of 20mM Octopamine (blue) or vehicle (gray). Green shading denotes the time of OA injection. Two-way repeated measures ANOVA finds a significant time-by-treatment interaction (F_(143,5863)_=1.651, p<0.0001). **(l-m)** Total sleep **(l)** and normalized activity counts **(m)** during 24h post-injection for groups shown in **(k)**. At least one distribution in **(l)** and **(m)** fail D’Agostino & Pearson test for Normality; Mann-Whitney tests find U= 125, p=0.0092 for **(l)** and U=110.5, p=0.0051 for **(m)**. For **(k-m)**, n= 21 vehicle control and 22 OA-injected flies.

*D. mojavensis* sleeps more than *D. melanogaster* and responds to prolonged waking with increased recovery sleep, indicating that this species may have an increased need for sleep relative to *D. melanogaster*. In their desert habitats, *D. mojavensis* are exposed to environmental stressors, including temperature variations and periods of sparse food and/or water availability. To measure sleep during desert-like temperature fluctuations, we exposed both *D. melanogaster and D. mojavensis* flies to daytime temperature ramps that held flies at 25°C overnight, then began to progressively increase the temperature across the first 6h of daytime to a peak of 35°C before reducing back to 25°C by lights-off at ZT12. While *D. mojavensis* maintained higher amounts of sleep than *D. melanogaster* across most of the day during these conditions **(Fig. 4a)**, both species showed a brief decrease in sleep when temperature peaked at 35°C at mid-day. As the temperature decreased afterwards, *D. melanogaster* briefly increased their sleep to comparable levels as the desert-adapted *D. mojavensis* subspecies. These results indicate that sleep in both species can be altered by variations in temperature, but that *D. mojavensis* flies retain elevated levels of daily sleep under naturalistic daytime temperature conditions.

**Figure 4.**
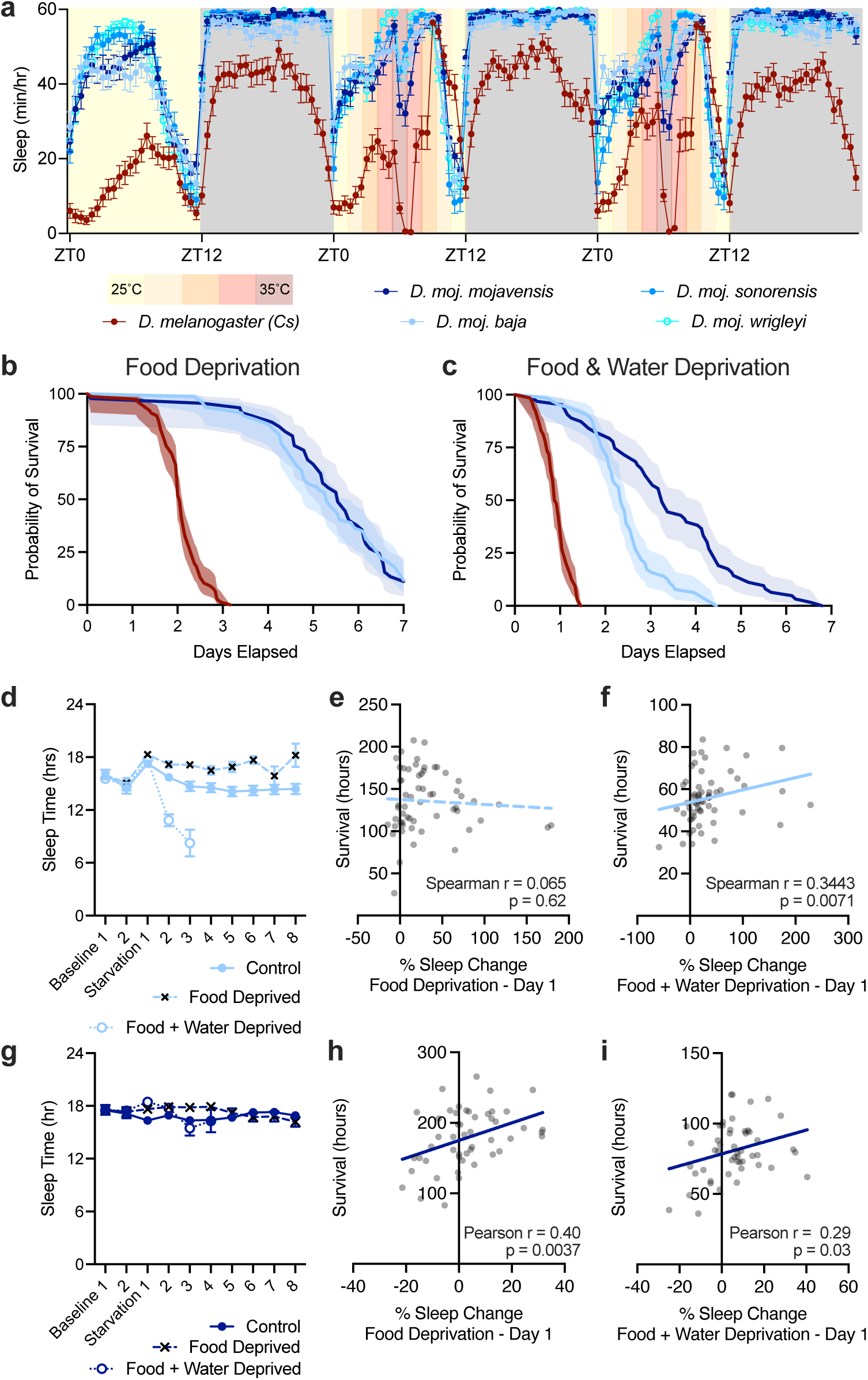
Sleep responses of *D. mojavensis* to nutrient deprivation correlate with survival time. **(a)** Sleep time courses for *D. melanogaster* (Cs; red) and *D. mojavensis* (blues) flies that were housed at 25°C for one baseline day, then exposed to a temperature ramp from 25°C to 35°C and back to 25°C during the daytime (ZT0-12). Two-way repeated measures ANOVA finds a significant time-by-strain interaction (F_(568, 26412)_=9.054, p<0.0001, n= 40 *Cs*, 39 *D. moj. moj*, 35 *D. moj. baja*, 35 *D. moj. sonorensis*, 42 *D. moj. wrigleyi*). **(b-c)** Survival times for *D. melanogaster* (*Cs*; red) and *D. mojavensis* (blues) females when housed on starvation agar **(b)** or dry tubes **(c)**. Mantel-Cox test finds significant effects for **(b)** (χ^2^=250.6, df=2, p<0.0001, n= 70 *Cs*, 45 *D. moj. moj.*, 77 *D. moj. baja*) and for **(c)** (χ^2^=232.6, df=2, p<0.0001, n= 64 *Cs,* 63 *D. moj. moj*, 64 *D. moj. baja*). **(d)** Daily sleep time for *D. moj. baja* flies during two days of baseline conditions that were then fed standard fly media (closed blue circles), 1% agar in water (black crosses), or dry tubes (open circles). Mixed-effects analysis finds significant effects of condition (p=0.0005) and time (p<0.0001), n = 63 control, 63 food deprived, and 64 food and water deprived flies at beginning of experiment. **(e-f)** Individual changes in sleep time on food deprivation day 1 **(e)** or food and water deprivation day 1 **(f)** plotted against survival during food deprivation for individual *D. moj. baja* females. At least one distribution in **(e)** and **(f)** failed the D’Agostino and Spearman test for Normality. A significant Spearman correlation was detected for food and water deprivation (**f**; r = 0.3443, p=0.0071, n=60 flies), but not for food deprivation (**e**; r= 0.065, p=0.62, n=60 flies). **(g)** Daily sleep time for *D. moj. moj.* flies during two days of baseline followed by feeding either standard fly media (closed circles), 1% agar in water (black crosses), or dry tubes (open circles). Mixed-effects analysis detected no effect of condition (p=0.7986), but a significant effect of time (p=0.0396), n = 56 control, 53 food deprived, 54 food and water deprived flies at the beginning of the experiment. **(h-i)** Changes in individual sleep time of *D. moj. moj.* on day one of food deprivation **(h)** or food and water deprivation **(i)** plotted against survival time. A significant Pearson correlation was found for both food deprivation (**h**; r=0.40, p<0.0037, n=50 flies) and food and water deprivation (**i**; r=0.29, p=0.03, n=50 flies).

To further test the functional relevance of heightened sleep pressure in desert-adapted flies, we also measured sleep and survival while flies were deprived of food alone or both food and water. Both Baja and Mojavensis subspecies of *D. mojavensis* survive longer than wild-type *D. melanogaster* when housed in glass tubes with non-nutritive agar media **(Fig. 4b)** or, as previously found, in empty, dry glass tubes **(Fig. 4c)**. While wild-type *D. melanogaster* suppress their sleep during food deprivation^46, 47^, *D. mojavensis* instead augment their sleep under these conditions. As shown in **Fig. 4d**, the sampled *D. moj. baja* female population significantly increased their total sleep time from day 2 of food deprivation, suppressing sleep only when lacking both food and water. *D. moj. Mojavensis* sleep responses to food deprivation are similarly increased for the first four days, after which sleep drops to control levels **(Fig. 4g, Extended Data Fig. 2a-b)**. Under food deprivation, both Baja and Mojavensis show increased consolidation of sleep, with fewer and longer sleep bouts as compared to controls **(Extended Data Fig. 2c-f)**. In contrast to Baja flies, the Mojavensis subspecies also shows increased (day 1) or control levels of sleep (days 2-4) when deprived of both food and water along with increased survival over Baja **(Fig. 4c, Fig. 4g)**.

Our results are consistent with the hypothesis that increased sleep during nutrient deprivation is associated with prolonged survival. Next, we tested this relationship at the individual level by plotting the change in sleep from baseline to starvation day 1 against the survival duration for each individual fly. In **Fig. 4e**, we find no significant relationship between sleep responses on day 1 of food deprivation and survival time for *D. moj. baja*. In contrast, we detect a significant correlation between relative change in sleep on experimental day 1 and survival time for individual *D. moj. baja* flies that are deprived of both food and water **(Fig. 4f)**. When we examined variation in the responses of individual *D. moj. mojavensis* flies, we also found a positive relationship between changes in sleep amount and survival duration when *D. moj. mojavensis* were either deprived of food **(Fig. 4h)** or of both food and water **(Fig. 4i)**. Our findings indicate that individual sleep responses during food and water deprivation (both subspecies) or for food deprivation (*D. moj. mojavensis*) are positively correlated with survival time. Interestingly, food-deprived *D. moj. baja* flies are hyperactive, with increased activity per time awake **(Extended Data Fig. 2g)**. Thus, though total wake time is decreased, total activity remains at control levels **(Extended Data Fig. 2h)**. In contrast, food-deprived *D. moj. mojavensis* flies display no hyperactivity, and total activity is reduced **(Extended Data 2i-j)**.

Preserved waking energy expenditure in food-deprived *D. moj. baja* might underlie the absence of a correlation between individual sleep increases and increased survival. Periods of adaptive sleep loss have been reported in several vertebrate species, especially in birds^48^ and marine mammals^49, 50^. During these periods, it is thought that animals can acutely cope with the costs that accumulate from sleep loss. Here, we find that *D. mojavensis* exhibits an opposing behavioral strategy: they chronically show elevated sleep time and consolidation, even during periods of insufficient food. This adaptive strategy is associated with a survival advantage in conditions of hunger or thirst. While additional work is required to identify and characterize functional advantages that are fulfilled by elevated levels of sleep, it is possible that hypersomnia may allow desert-adapted *D. mojavensis* to maintain efficient energy usage^51–53^, clearance of metabolic waste^54, 55^, or management of oxidative stress^56, 57^. Alternatively, the increased need for sleep in *D. mojavensis* could offset costs of physiological adaptations made by desert-adapted flies that allow them to feed on cactus diets or to thrive in the desert environment^16, 58–63^. Interestingly, another recent study found that other *Drosophila* species exhibit a range of homeostatic responses to sleep loss^64^, indicating that broad studies of *Drosophila* evolution could uncover species-specific adaptations in sleep need or function. Our characterization of increased sleep time and need in stress-resilient *D. mojavensis* provide a novel model species to examine the adaptive advantage(s) of elevated sleep and to investigate the evolution of sleep regulatory mechanisms across related species.

## Acknowledgements

We thank all members of the Donlea lab for many helpful discussions and technical advice. Special thanks to Dr. David Krantz (UCLA) for technical assistance and feedback. We thank Dr. Luciano Matzkin at Arizona State University for sharing wild-caught *D. mojavensis (baja, mojavensis, sonorensis, and wrigleyi).* Many thanks to Dr. Orkun Akin (UCLA) for kindly sharing access to his lab’s confocal microscope and to Dr. Chris Vecsey (Colgate University) for sharing unpublished updates to SCAMP sleep analysis scripts. CN is supported by the NIH IRACDA program at UCLA NIH K12GM106996. SW is funded by support from NSF-IOS 2016188. JMD is funded by support from NIH R01NS105967 and R01NS119905.

## Author Contributions

J.Y., C.N., K.L, P.H., C.J., M.R., D.T., X.H., and J.M.D performed the experiments and/or analyzed data. G.E. and J.P.W. provided consultations, completed LC-MS experiments, and analyzed the data. S.W. and J.M.D. initially discussed and designed the project. J.M.D. supervised the research. J.Y., C.N., and J.M.D. integrated the data, interpreted the results, and wrote the manuscript. All authors discussed the results and commented on the manuscript.

## Competing Interests

The authors declare no competing interests.

## Methods

### Fly Rearing and Stocks

Fly stocks were cultured on standard cornmeal molasses media (per 1L H_2_O: 12 g agar, 29 g Red Star yeast, 71 g cornmeal, 92 g molasses, 16mL methyl paraben 10% in EtOH, 10mL propionic acid 50% in H_2_O) at 25°C with 60% relative humidity and entrained to a daily 12h light, 12h dark schedule. Experiments with Banana-Opuntia media used a recipe from the National *Drosophila* Species Stock Center (NDSSC; Cornell University): per 1L H_2_O: 14.16g agar, 27.5 g yeast, 2.23g methyl paraben, 137.5g blended bananas, 95g Karo Syrup, 30g Liquid Malt Extract, 22.33g 100% EtOH, 2.125g powdered opuntia cactus.

*Canton-S* were provided by Dr. Gero Miesenböck (University of Oxford) and *Pcf* were shared by Dr. Mark Frye (UCLA). Primary stocks of *D. moj. mojavensis, D. moj. baja, D. moj. wrigleyi*, and *D. moj. sonorensis* were a gift from Dr. Luciano Matzkin (University of Arizona), and additional stocks of *D. moj. mojavensis* and *D. moj. baja* were shared by Dr. Paul Garrity (Brandeis University). *D. arizonae* flies (SKU: 15081-1271.36) were ordered from the NDSSC. Wild caught *D. melanogaster* descended from a single pair of flies trapped in Los Angeles, CA in spring, 2020.

### Behavior

4-8 day old female flies were housed individually in borosilicate glass tubes (65mm length, 5mm diameter) containing fly food coated with paraffin wax at one end and a foam plug in the other. Locomotor activity was recorded using DAM5M or DAM5H multibeam *Drosophila* Activity Monitors from Trikinetics Inc. (Waltham MA, USA) and sleep was analyzed in Matlab (MathWorks Inc) with the SCAMP script package^65^. Locomotor activity was measured as the number of movements between beams per one-minute bins. Periods of sleep were defined by at least 5 minutes with no change in position within the multibeam activity monitors.

### Sleep Deprivation and Arousability

Sleep deprivations were performed mechanically by mounting DAM5M activity monitors onto platform vortexers (VWR 58816-115). Individual tubes were plugged with food at one end and 3D-printed PLA plastic caps at the other. Monitors were vortexed at an intensity of 2.5g for 3-second pulses every minute through the duration of the 12-hour dark period. Arousability was tested in a darkened incubator with 60 seconds of blue light (luminance 0.048 Lv) every hour for 24 hours following sleep deprivation.

### Food- and Water- Deprivation Assays

All flies were put in DAM5H activity monitors on standard food for baseline recording. After 2-3 days, control flies were transferred to tubes containing fresh food, food-deprived flies to tubes containing a 1% agar gel, and food-and-water-deprived flies to empty tubes plugged with foam at both ends. Flies immobile for at least 24 hours were defined as dead and data subsequent to their last full day alive was removed from analysis.

### Pharmacological Microinjections

4-8 day old female flies were loaded into behavior tubes and monitored in DAM5M Activity Monitors to obtain baseline sleep and locomotor activity under 12h light: 12h dark (25°C). After 1-2 days of baseline in DAM5M monitors, flies housed in borosilicate tubes were placed on ice for anesthetization prior to injection using Drummond Nanoject II. For injection of exogenous neuromodulators, the anteriormost ocelli of *D. mojavensis baja* were injected with 18.4nl of 20mg/mL of Octopamine (Sigma-Aldrich, Catalog # O0250). For each round of injections, new OA is solubilized using Schneider’s *Drosophila* Medium with L-Glutamine (Genesee Scientific, Catalog # 25-515). Following each individual injection, flies are returned back into individual borosilicate tubes, and placed in respective DAM5M Activity Monitors to continue sleep and activity surveillance for >48h.

### Immunohistochemistry

Female *D. melanogaster* and *D. mojavensis* were reared in 12h light:12h dark schedule at 25° C in normal fly food. Individual fly brains were dissected 5-7 days post-eclosion between a ZT0- ZT3 window to minimize time-of-day variation to antibody targets. All dissections, antibody staining, and preparation for imaging were carried out in the exact same manner to minimize variability when comparing between species. Flies are anesthetized using ice. Brains were dissected in chilled 1X PBS then placed in 4.0% paraformaldehyde/1X PBS (PFA) for 30 mins. in room temperature on a benchtop rotator. PFA from brains were removed by washing with 1.0% Triton-X in 1X PBS 3 times for 10 mins. each. Once brains were free of PFA, the brains were placed in 1x Sodium Citrate (10mM, pH=6.0, 15 mins. at 80° C) for antigen retrieval.

Brains were then placed in a blocking buffer (5.0% normal goat serum in 0.5% Triton-X/1X PBS) and incubated at room temperature for 1.5h on a rotator. Brains were incubated with one the following primary antibodies (diluted using blocking buffer): 1:1000 Mouse anti-PDF (Developmental Studies Hybridoma Bank). Primary antibodies were incubated for two days in 4°C. After incubation, brains we’re washed using 0.5% Triton-X in 1X PBS five times, 10 mins. each. Fly brains were then incubated in AlexaFluor secondary antibodies (1:1000 Goat anti-Mouse AlexaFluor 633nm; Molecular Probes) overnight at 4°C. Brains were washed using 0.5% Triton-X in 1X PBS five times, 10 mins. After washing, brains were mounted on glass slides in Vectashield mounting media, sealed with a coverslip and nail polish. Brains were imaged using a Zeiss LSM 880 laser scanning confocal microscope using a z-slice thickness of 1um and saved as CZI files. Maximum intensity projections were created from CZI files using FIJI/ImageJ (https://imagej.net/software/fiji/)^66^.

### Neurochemical Quantifications

#### Sample preparation protocol

The samples containing fly brains stored at -80°C are treated with 99.9/1 Water/Formic Acid. An internal standard (IS) of each targeted compound was added to every sample to account for compound loss during sample processing. The samples are vortexed, homogenized for 30 sec in a bead beater using 2.0 mm zirconia beads, and centrifuged at 16.000xg for 5 min. The supernatant is transferred to new microcentrifuge test tubes and dried in a vacuum concentrator. The samples are reconstituted in 40 µl of water, vortexed, and centrifuged. The supernatant is transferred to HPLC vials and 10 µl is injected to an HPLC -triple quadrupole mass spectrometer system for analysis.

#### Liquid Chromatography-Tandem Mass Spectrometry LC-MS

A targeted LC-MS/MS assay was developed for each compound using the multiple reaction monitoring (MRM) acquisition method on a triple quadrupole mass spectrometer (6460, Agilent Technologies) coupled to an HPLC system (1290 Infinity, Agilent Technologies) with an analytical reversed phase column (GL Sciences, Phenyl 2 µm 150 x 2.1 mm UP). The HPLC method utilized a mobile phase constituted of solvent A (100/0.1, v/v, Water/Formic Acid) and solvent B (100/0.1, v/v, Acetonitrile/Formic Acid) and a gradient was used for the elution of the compounds (min/%B: 0/0, 10/0, 25/75, 27/0, 35/0). The mass spectrometer was operated in positive ion mode and fragment ions originating from each compound was monitored at specific LC retention times to ensure specificity and accurate quantification in the complex biological samples (Octopamine OA 159-136, Histamine HA 112-95, Dopamine DA 154-137, Serotonin 5HT 177-160). The standard curve was made by plotting the known concentration for each analyte of interest (CDN Isotopes) against the ratio of measured chromatographic peak areas corresponding to the analyte over that of the labeled standards. The trendline equation was then used to calculate the absolute concentrations of each compound in fly brain tissue.

### Statistical Analysis

Statistical tests were completed as described in the figure legends using Prism 9 (GraphPad Software, Boston MA, USA). Statistical comparisons primarily consist of one-or two-way ANOVAs followed by pairwise Holm-Sidak’s multiple comparisons test when experiments include at least three experimental groups or two-tailed Student’s T-test for experiments that include two groups. All data figures pool individual data points from at least two independent replicates.

---

**Extended Data Figure 1.**
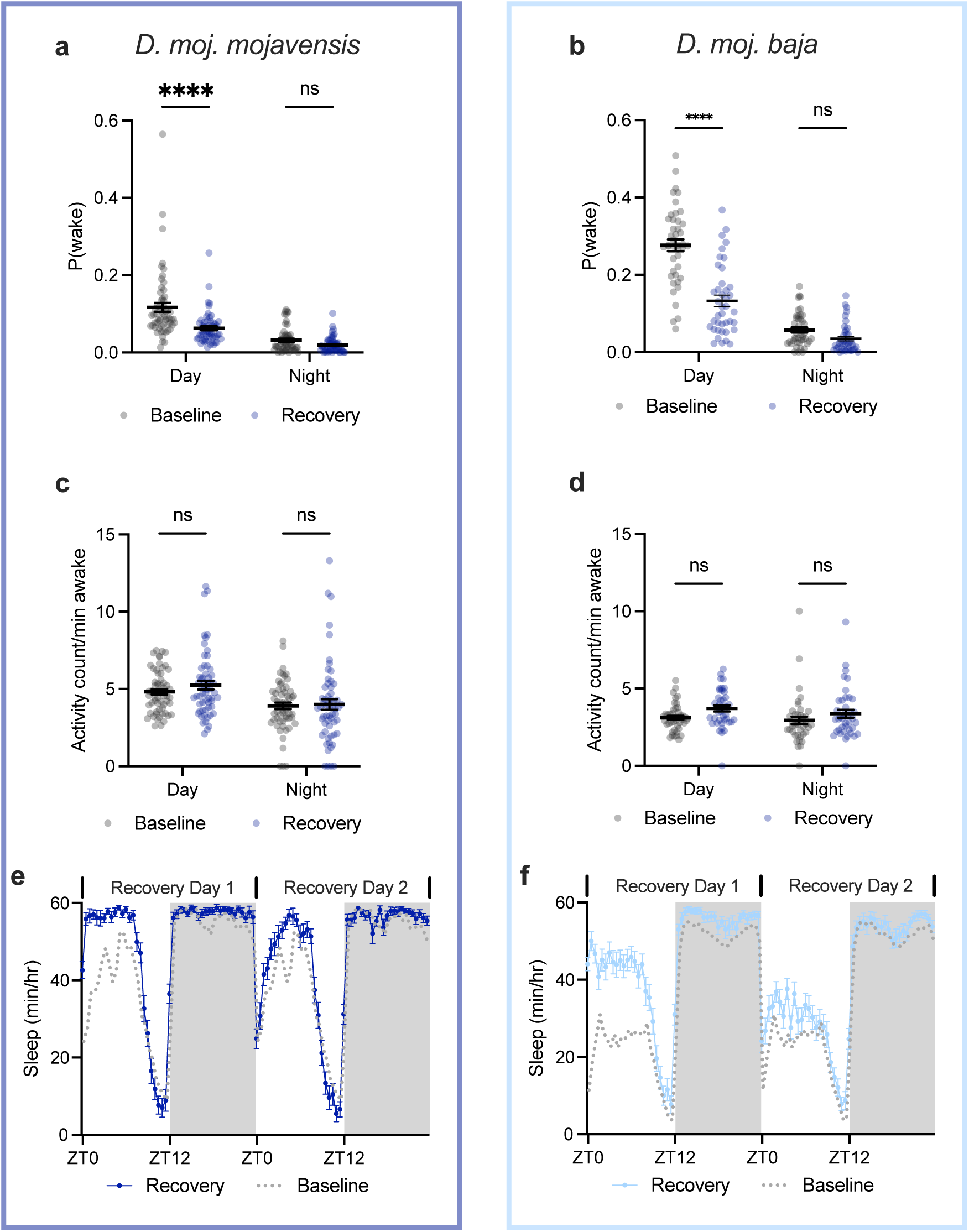
Sleep and activity parameters following sleep deprivation. **(a-b)** P(wake) for *D. moj. mojavensis* **(a)** and *D. moj. baja* **(b)** during baseline (gray) and first recovery day (blue) after overnight vortex sleep deprivation. Two-way repeated measures ANOVAs find significant day-by-time interaction for **(a)** F_(1,58)_=17.08, p=0.0001, n=59 flies and **(b)** F_(1,36)_=39.14, p<0.0001, n=40 flies/group. **(c-d)** Activity counts/waking minute for *D. moj. mojavensis* **(c)** and *D. moj. baja* **(d)** during baseline (gray) and first recovery day after vortex sleep deprivation (blue). Two-way repeated measures ANOVA find no significant main effect of day for **(c)** F_(1,58)_=1.061, p=0.31, n=59 flies, but do find a significant effect of day for **(d)** F_(1,42)_=5.509, p=0.0237, n=40 flies/group. **(e-f)** Sleep timecourses during two days of recovery following overnight sleep deprivation in *D. moj. mojavensis* **(e)** or *D. moj. baja* **(f)**. Sleep traces from 24h of baseline replotted in gray, recovery days shown in blues.

**Extended Data Figure 2.**
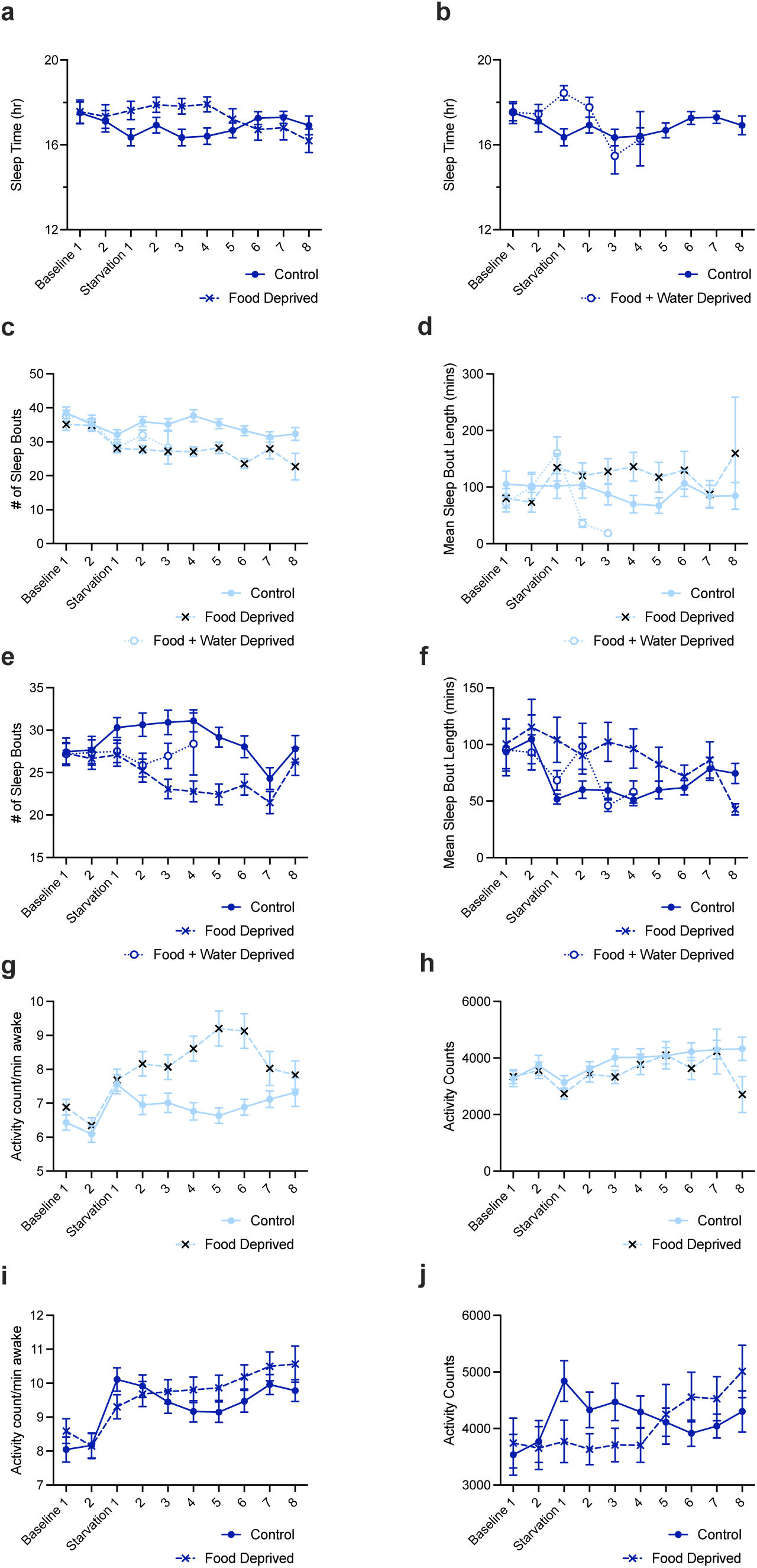
Sleep and activity parameters during food deprivation. **(a-b)** Daily sleep totals for *D. moj. mojavensis* flies that were fed (**a-b**, filled circles), food deprived (**a**, crosses), or food and water deprived (**b**, open circles). Data are replotted from Fig. 4g for visualization, See legend of Fig. 4g for mixed effects results, n= 56 control, 53 food deprived, 54 food and water deprived. **(c-d)** Sleep bout number **(c)** and sleep bout lengths **(d)** for groups of *D. moj. baja* shown in Fig. 4d. Mixed effects analysis effect of condition for **(c)** F_(2,189)_=9.459, p=0.0001 and for **(d)** F_(2,189)_=0.7335, p=0.4816, n=63 control, 63 food deprived and 64 food and water deprived flies/group. **(e-f)** Sleep bout number **(e)** and sleep bout lengths **(f)** for groups of *D. moj. mojavensis* shown in Fig. 4g. Mixed effects analysis effect of condition for **(e)** F_(2,163)_=5.511, p=0.0048 and for **(f)** F_(2,163)_=2.922, p=0.0566, n= 56 control, 53 food deprived, 54 food and water deprived flies. **(g-h)** Mean counts/waking minute **(g)** and total activity counts **(h)** for *D. moj. baja* flies shown in Fig. 4d. Mixed effects analysis effect of condition for **(g)** F_(2,189)_=16.39, p<0.0001 and for **(h)** F_(2,189)_=17.36, p<0.0001, n=63 flies/group. **(i-j)** Mean counts/waking minute **(i)** and total activity counts **(j)** for *D. moj. mojavensis* flies shown in Fig. 4g. Mixed effects analysis effect of condition for **(g)** F_(2,163)_=0.2880, p=0.7501 and for **(h)** F_(2,163)_=0.01571, p=0.98, n= 56 control and 53 food deprived flies.

## Notes

### Competing Interest Statement

The authors have declared no competing interest.

